# Optimal Point Process Filtering for Birth-Death Model Estimation

**DOI:** 10.1101/165712

**Authors:** Kris V Parag, Oliver G Pybus

## Abstract

The discrete space, continuous time birth-death model is a key process for describing phylogenies in the absence of coalescent approximations. Extensively used in macroevolution for analysing diversification, and in epidemiology for estimating viral dynamics, the birth-death process (BDP) is an important null model for inferring the parameters of reconstructed phylogenies. In this paper we show how optimal, point process (Snyder) filtering techniques can be used for parametric inference on BDPs. Specifically, we introduce the Bayesian Snyder filter (SF) to estimate birth and death rate parameters, given a reconstructed phylogeny. Our estimation procedure makes use of the equivalent Markov birth process description for a reconstructed birth-death phylogeny (Nee *et al*, 1994). We first analyse the popular constant rate BDP and show that our method gives results consistent with previous work. Among these results is an analytic solution to the special case of the Yule-Furry model. We also find an equivalence between the SF Poisson likelihood and two standard conditioned birth-death model likelihoods. We then generalise our estimation problem to BDPs with time varying rates and numerically solve the SF for two illustrative cases. Our results compare well with a recent Markov chain Monte Carlo method by Hohna *et al* (2016) and we numericaly show that both methods are solving the same likelihood functions. Lastly we apply the SF to a model selection problem on empirical data. We use the Australian Agamid dataset and predict the same relative model fit as that of the original maximum likelihood technique developed and used by Rabosky (2006) for this dataset. While several capable parametric and non-parametric birth-death estimators already exist, ours is the first to take the Nee *et al* approach, and directly computes the posterior distribution of the parameters. The SF makes no approximations, beyond those required for parameter space discretisation and numerical integration, and is mean square error optimal. It is deterministic, easily implementable and flexible. We think SFs present a promising alternative parametric BDP inference engine.

## 1. Introduction

Real populations are discrete, denumerable and undergo random fluctuations in size due to births (speciations or branching events) and deaths (extinctions). Consequently, they are most accurately described by stochastic, continuous time birth-death branching processes (BDPs) [1]. In its classical form (known as constant rate BDP), if the discrete integer state of the BDP is *l*(*t*) at time *t* then the model proceeds in forward time by allowing one of two nearest neighbour transitions. Either a birth occurs to state *l*(*t*) + 1 with rate *λl*(*t*) or a death occurs to state *l*(*t*) – 1 with rate *μl*(*t*). For our purposes *l*(*t*) will be called the true number of lineages or taxa. The pair of non-negative constants (*λ, μ*) are called the per lineage birth and death rates. Births or deaths of any lineage occur independently of any other and at any time only one transition is possible. Assuming *l*(0) = 1 and allowing the BDP to progress with time results in a rooted binary tree (phylogeny) containing both extant and extinct members, that is a useful null model for molecular phylogenies [2]. To ensure the tree survives to the present, *T* > 0, this process is often conditioned so that *l*(*T*) ≥ 1. This is representative of the idea that if a BDP did not survive we would have no way of observing it. Practically, it is only possible to observe dead lineages or extinct taxa if fossils are available. In our work, we focus on the BDP in the absence of fossil data.

If the BDP tree is pruned such that only lineages with descendants at *T* remain then this is called the reconstructed birth-death process (rBDP) and corresponds to an ideal observable phylogeny [2]. We denote the number of lineages in the reconstructed tree at time *t* by the counting process 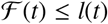. 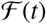 for 0 ≥ *t* ≥ *T* is a non-decreasing integer function that describes the observable data (it counts reconstructed branching times). Harvey *et al* [3] showed that even though the rBDP does not have explicit information on extinct lineages it still contains signal for estimating both the per lineage birth and death rates (*λ, μ*). Estimating these rates given 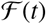, for the constant rate BDP and more generalised BDPs, is of central importance to macroevolution, ecology and phylodynamics, since it allows us to fundamentally link the properties of organisms to their environment [4].

In particular, inference solutions for BDPs with time-varying rates, density dependent rates, trait dependent rates, incomplete sampling schemes, phylogeographic and multi-type behaviours have led to many insights ranging from understanding the diversification behaviour in the animal kingdom to the space-time dynamics of viral epidemics [5] [6] [7] [8] [9] [10]. Achieving these insights is, however, not an easy task as the loss of information (death events) in going from a true BDP (complete phylogeny) to an rBDP (observable phylogeny) leads to several difficulties. For example many combinations of birth and death rates can produce the same rBDP which can cloud the true diversification process and result in inaccurate or biased rate estimates [11] [4]. Moreover, many standard inference methods may struggle to infer non-zero death rates from real data despite known extinction events [4] [12]. This is often attributed to model assumptions being implicitly violated or an inability to properly describe the diversification scenario presented by the data [13]. Additionally, validating these BDP estimates and insights against those achievable by other competing phylogenetic models such as the coalescent [14] is also of interest [15] [6] [16]. Since it is believed that many of these points can be addressed or resolved with more sophisticated inference schemes that account for more complex rate variation settings [13], BDP inference remains an important, non-trivial and current field of research.

We focus on inference for BDPs with time-varying birth and death rates, denoted (*λ*(*t*), *μ*(*t*)). Such temporal variance is often attributed to external influences such as abiotic and biotic environmental changes [4] [13]. In particular, we introduce a parametric Bayesian estimation method from control and electrical engineering which we refer to as the Snyder filter (SF) [17]. We apply it to infer *λ*(*t*) and *μ*(*t*) given the rBDP, and treat the constant rate BDP as an important special case. This application rests on having a description of the rate of producing new observable events according to the rBDP. We previously showed how the SF could be adapted for inference on the coalescent process, the most popular competing phylogenetic model to the BDP [18]. While we do not discuss the performance of BDPs relative to coalescent models, we do comment on when their inference problems are deemed mathematically similar according to the SF approach.

Several methods already exist for inferring time-varying BDP rates from rBDPs. Nee *et al* [2] initiated the investigation of rBDPs in 1994 by writing down, but not optimising, likelihood functions based on multiplying the densities of waiting times for events in the rBDP. Since then explicit likelihood function approaches have dominated the field [13] with several methods based on different: constructions of probability densities, conditioning criteria for the BDP, optimisation scheme choices and on whether a maximum likelihood or Bayesian viewpoint is emphasised [9]. A detailed look at the various likelihood based inference methods in the literature for time-varying and even more generalised BDPs can be found in Morlon *et al* [13]. The most powerful among these methods tend to use Markov chain Monte Carlo (MCMC) sampling to optimise likelihood functions. This allows one to accommodate complex BDP features such as sampling schemes, clade variation or genealogical uncertainty, usually at the expense of analytic exactness and methodological simplicity [6] [7].

Existing non-likelihood based methods usually involve summary statistics or optimisation to lineage through time plots (LTTs) [19] [13] [20]. In our notation LTTs are plots of the number of reconstructed lineages 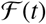 against *t*. Such methods often do not account for the aforementioned complex BDP features and possess less power. However, they are still popular since they are simple and easy to use. The SF technique we introduce in this paper functionally fits between the likelihood and non-likelihood approaches. We focus on achieving an analytical or exact solution as much as possible while maintaining ease of use and avoiding optimisation and sampling procedures like MCMC. We do not yet account for complexity like genealogical uncertainty or non-uniform sampling schemes but we do submit a method with the potential flexibility to accommodate such features in the future. Methodologically, the SF has very different mechanics from other existing techniques since it simply involves solving coupled linear ordinary differential equations with discontinuous updates at branching times. The result is a direct computation of the posterior distribution of the parameters of interest given the observable rBDP.

We start by defining the SF and its equations in the Methods section before describing how it can be applied to BDPs which are conditioned to survive until some arbitrary present time, *T*. We also explain how to incorporate homochronous sampling but do not implement this addition since it does not alter the inference problem. Consequently, all our work is based on completely sampled BDPs. In the Results and Analysis section we first numerically solve the constant rate BDP Snyder equations and then compare results against both the true values and a least squares method by Paradis [20]. Further, we provide analytic Snyder solutions for the Yule-Furry process and make some comparisons to the Kingman coalescent. We then illustrate that the Poisson type likelihood solved by the SF is equivalent to the standard likelihood of a BDP conditioned on survival. This equivalence holds for trees starting at either the root or crown (most recent common ancestor). We extend our analysis to time-varying birth and death rates and test the filter on two simulated models from the literature. We compare our results with the true value and a modern, recent Markov chain Monte Carlo (MCMC) based method by Hohna *et al* [8] [21]. We not only find that estimates correspond but also show that the likelihoods evaluated by both methods are well matched. Finally we show how the SF can be used for model selection on empirical data. We run the filter on the Australian Agamid dataset and compare our results with those from the maximum likelihood method developed by Rabosky [22]. Rabosky originally tested his method on this Agamid dataset. In the Discussion we comment on the ability and potential of the SF as a useful inference technique for BDPs and note that it can easily incorporate extra functionality such as density dependence.

## 2. Methods

### 2.1. Optimal Snyder Filtering

Consider a doubly stochastic Poisson process (DSPP), 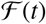, at time *t* ≥ 0. 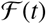 is a non-decreasing integer valued process that counts the number of points at t. Its instantaneous intensity, 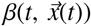, is stochastic and on the space of non-negative real numbers. We want to infer the vector information process 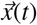 (section 2.2 will show how 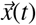 encodes the parameters of a BDP) given past observations of the DSPP. We denote the observed process as 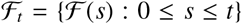. Snyder derived a filter that solved the problem of optimally inferring 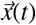 given 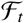 and the basic statistics of 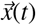 [17]. We call this the SF. It is an exact, Bayesian inference method that generates the informed posterior, 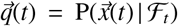, by solving a set of non-linear differential equations on the probability distribution of 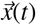, sequentially with time over 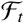. The SF is ‘causal’ because the inferred posterior at time *t* only depends on observations up to *t*. It is ‘exact’ because it computes this joint posterior directly, without approximating either the observation process, 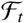, or the information process, 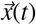. The only approximations in this method are due to the standard inaccuracies in numerically integrating its differential equations and in representing distributions discretely. In some cases, these differential equations are analytically solvable and there are no approximations.

The SF is quite ‘flexible’ since it applies to any Markov information process that has dynamics describable by: 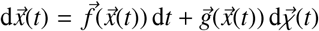. Here 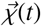 is a martingale with independent increments and 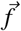, 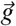 are arbitrary vector functions of choice (see [23] for details). The filter is also valid if the DSPP intensity is generalised to depend on the observed process as 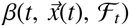. This makes the DSPP self-exciting [23]. The SF framework is therefore very general. In section 2.2 we will show how estimation with the reconstructed BDP can be cast as a Snyder inference problem. We focus on inference problems with 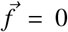 and 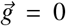. This means our information process is just a vector of random variables 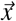. We also restrict the type of self-exciting dependence in the DSPP rate to be Markovian (0-memory). This means that the DSPP intensity is now 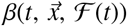 instead of 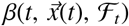. Under these conditions the SF, can be transformed into a set of linear differential equations on an un-normalised distribution *q*\*(*t*). This is then normalised afterwards to 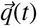 [24]. The resulting SF is described by equations 1–3 [23] [24]. 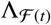 is a diagonal matrix called the rate matrix. It has entries for each possible value of 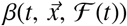 at any given *t* due to the possible values of 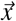 and 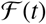. Let an arbitrary value of 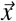 be *ϵ* and denote the corresponding normalised and unnormalised probabilities as 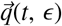 and 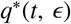. The complete posterior distribution is then 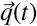 while 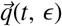 is the single value corresponding to *ϵ*. Assume we have observed an event stream over 0 ≤ *t* ≤ *T*. If the first event time is *t* = *τ*_1_ then 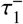 and 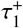 are infinitesimally before and after that event time. The initial condition for the differential equations is the prior: 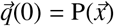. The following equations then describe the dynamics of 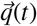 with time until 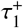.

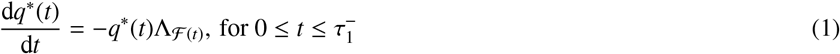

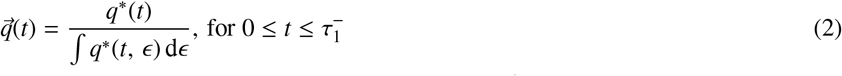

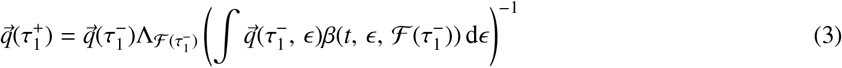

From 0 to 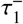 the state probabilities undergo a continuous exponential decaying trajectory (equation 1). At this point an event is observed and a discontinuous update of the posterior occurs (equation 3). The updated posterior 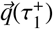 is then used as a new initial condition and the equations solved again until the next event (over 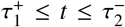). This procedure repeats until we obtain 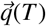). Note that the integrals enumerate every possible value of 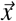. More information on the SF and some of its biological applications can be found in [17] [23] [25] and [26] and [18].

We will show that BDP inference falls within the SF framework and derive the appropriate rate matrix 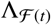. Our posterior of interest, 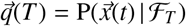 uses all of the observed data 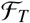 and yields the minimum mean squared error (MMSE) estimator of any given function of the parameters, 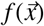. This is denoted 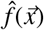 and defined:

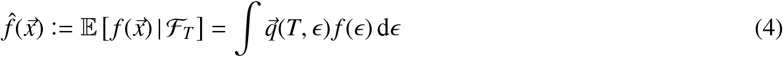

The estimator 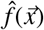 is also called the conditional mean [23]. The integrals are taken over the parameter vector space. The SF is optimal because it directly calculates the posterior, 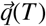, that achieves the MMSE.

### 2.2. Birth-Death Process Inference

We consider a BDP with time-varying but not lineage dependent rates. We denote the per lineage birth and death rates as *λ*(*t*) and *μ*(*t*) respectively and the number of true lineages or taxa at time *t* by *l*(*t*). Due to death events (extinctions) *l*(*t*) is unobservable. We assume that the BDP survives until some time *T* and denote the number of extant lineages at this time *l*(*T*) = *n*. If the true birth-death tree is pruned such that only lineages with descendants at T remain then this is called the reconstructed process (rBDP) and corresponds to an ideal observable phylogeny [2]. Let the number of lineages on the rBDP at time *t* be 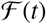 with 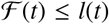 and boundary conditions 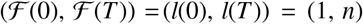. Unlike *l*(*t*) which rises and falls with time, 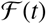 is a non-decreasing integer valued function. If we denote the time of the *k*^th^ birth event in the rBDP as *c_k_* for 1 ≤ *k* ≤ *n* – 1 then the observed process is: 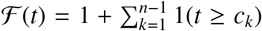. Here 1(:) is an indicator function. As is common in the literature we define the negative total diversification, *α*(*t, τ*) and the probability that a single lineage at time *t* survives until *T*, *P*(*t, T*), as follows. These equations were originally derived by Kendall [27].

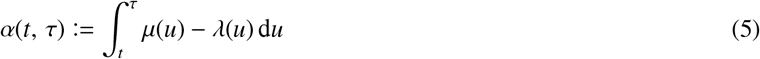

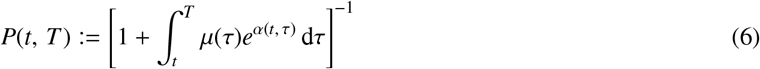

Nee *et al* [2] showed that the rBDP is actually a generalised pure birth process [27] with birth rate, *β*(*t*) defined below. This rate contains information about deaths through the *P*(*t, T*) function. Note that when *μ*(*t*) = 0 over 0 ≤ *t* ≤ *T* then *P*(*t, T*) = 1 and 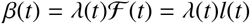 and the rBDP contains all the information of the BDP.

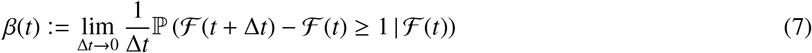

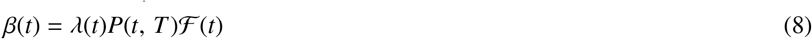

In this form the rBDP rate is amenable to SF inference. This is because the counting statistics of a self-exciting point process with 0-memory are identical to those of pure birth process with a population dependent birth rate [23]. We assume that the BDP can be described with *p* parameters: 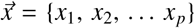 and that this set is partitioned so that the birth and death rates are parametrised as 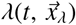 and 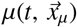. Note that 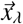 and 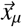 may have common parameters but together they must span all of 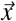. For notational brevity we will not explicitly refer to 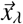 or 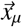 and continue writing *λ*(*t*) and *μ*(*t*). The inference problem on the rBDP therefore conforms to that of the self-exciting DSPPs mentioned in section 2.1. Our rate of interest is explicitly 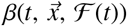 and we want to compute 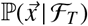.

We assume complete homochronous sampling so that all *n* taxa or lineages observable in the rBDP are sampled at the present time *T* with probability 1. Incomplete homochronous sampling would involve sampling each extant lineage with probability *ν* < 1. It is well documented that estimates change based on how the taxa are sampled [5]. The impact of sampling measures on inference is a problem that is currently outside the scope of our work. However, in Appendix A we explain how incomplete homochronous sampling can be easily handled by the current SF framework since it simply alters the form of *β*(*t*).

We now explain the numerical implementation of the filter for BDP inference. Let the vector of *p* random variables (parameters) to be estimated, 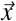 be such that the distribution of the *i*^th^ random variable can be described on a domain of *m_i_* points. The parameter vector is then on a joint Cartesian grid of 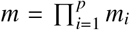 possible values and so there are *m* possible sets of values for 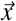. We denote some arbitrary value from this set of 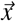 as *ϵ*. The SF solves a differential equation for the joint probability mass across all *ϵ*. Consequently, the filter has dimension *m* and the prior 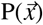 and posterior 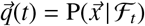 have *m* elements. The rate matrix, 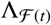 is then an *m* × *m* diagonal matrix with entries at each time *t* given by enumerating 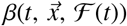 across the possible *ϵ*. The integrals in equations 1–3 are across the elements of the relevant vectors or matrices and solved along the observed stream 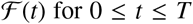. Note that at the rBPD event times (at which equation 3 is solved): 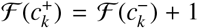. The only approximations of the Snyder inference method are inherited from the tolerances in performing standard numerical integration and the quantisation errors in discretising the parameter space. The MMSE estimates and the joint posterior 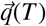 form the core of the results that we present in sections 3.1 – 3.5. We take estimates at *T* because this makes use of all the information that exists in the event time series. More information on the numerical implementation, accuracy and complexity of the SF as applied to phylodynamic problems can be found in Parag and Pybus [18].

## 3. Results and Analysis

We apply the SF to a series of birth-death inference problems. We start by estimating the constant rate BDP. We compare our results with known true parameter values and with a least squares method by Paradis [20]. We then provide an analytic solution to the Yule-Furry process (a constant rate BDP with zero death rate) and make comparisons to the Kingman coalescent. We also present an inhomogeneous SF solution which leads to a Snyder likelihood. We show numerically that this is identical to two of the well known, standard likelihoods for the constant rate BDP. We then extend our analysis to BDPs with time varying birth and death rates. These present a notably more difficult inference problem. We numerically solve filters for two example models from the literature and compare results, both in terms of sampled marginal posteriors and computed parameter likelihoods, with a recent fixed tree MCMC method developed by Hohna [8]. Having verified the SF on simulated models we extend our investigations to the empirical Australian Agamid dataset. We use the SF to perform model selection according to a simple least squares metric and observe the similarity between our estimates and those of a maximum likelihood method by Rabosky [22] that was originally tested on this dataset.

### 3.1. Constant Rate Birth-Death Estimation

The constant rate BDP with complete homochronous sampling is considered. The parameters to be estimated are set as *x*_1_ = *λ* and *x*_2_ = *μ* or equivalently 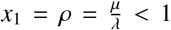 and *x*_2_ = *σ* = *λ* – *μ* > 0. The second parametrisation is standard in the literature [5]. For the constant rate, completely sampled BDP, *P*(*t, T*) has the explicit expression given in equation 9. We can then parametrise *β*(*t*) in terms of *σ* and *ρ* as in equation 10. We note in Appendix A, that solutions to this completely sampled problem are directly applicable to the incompletely sampled one.

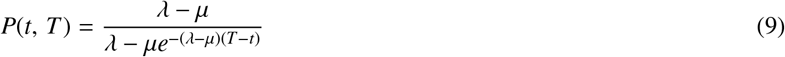

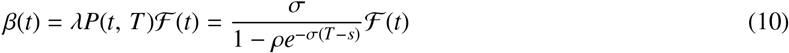

We start our BDP investigation by simulating a true constant rate rBDP under known parameter values. We used the efficient BDP sampling algorithms of [28] and [5] with re-parametrised inverse distributions given below. Here the uniform random variables 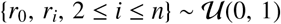 and we assume *n* extant lineages.

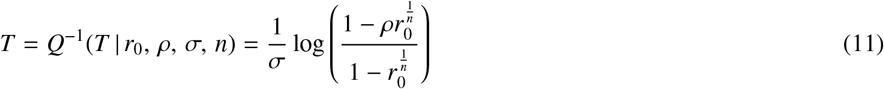

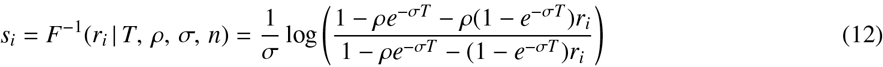

We obtain a tree depth *T* using equation 11 and then a series of bifurcation times *si* from expression 12. The *s_i_* represent the information carrying data, since we can map the entire reconstructed tree from these speciation times. In the original algorithm time is labelled going backward so that the present is time 0. We work in forward time from 0 to T and so construct the dataset *S* = sort{*T* – *s_i_*, 1 ≤ *i* ≤ *n*} where sort{*X*} sorts *X* in ascending order. The birth events are such that *c*_*n*–1_ = *T* with *c_k_* for 1 ≤ *k* ≤ *n* – 1 as the (*k* + 1)^th^ element of *S*. The observed process 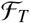 is then the counting process of these events and the *c_k_* are the times at which we perform the Snyder updates of equation 3.

We numerically integrate the SF (equations 1–3) using the Markov birth process description of [2] with the rate function *β*(*t*) from equation 10. We use the (*σ, ρ*) parametrisation since it is numerically easier to define priors under this pair. We then convert our results to the classical (*λ, μ*) for analysis. We obtained conditional mean estimates for these parameters over 10^4^ replications and plotted them in the histograms of Figure 1a at a high and low *μ* setting. The percentage relative MMSE is obtained for parameter *x_i_* according to 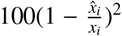. In all cases the Snyder estimates seem to cover the true values well. In general *μ* appears to be more difficult to estimate than *λ* and estimation accuracy improves with reduced *μ*. Lastly, when *μ* is high it seems that there is a bias toward underestimating its value. These are all well known trends in the literature which are now confirmed by the SF [29].

**Figure 1:**
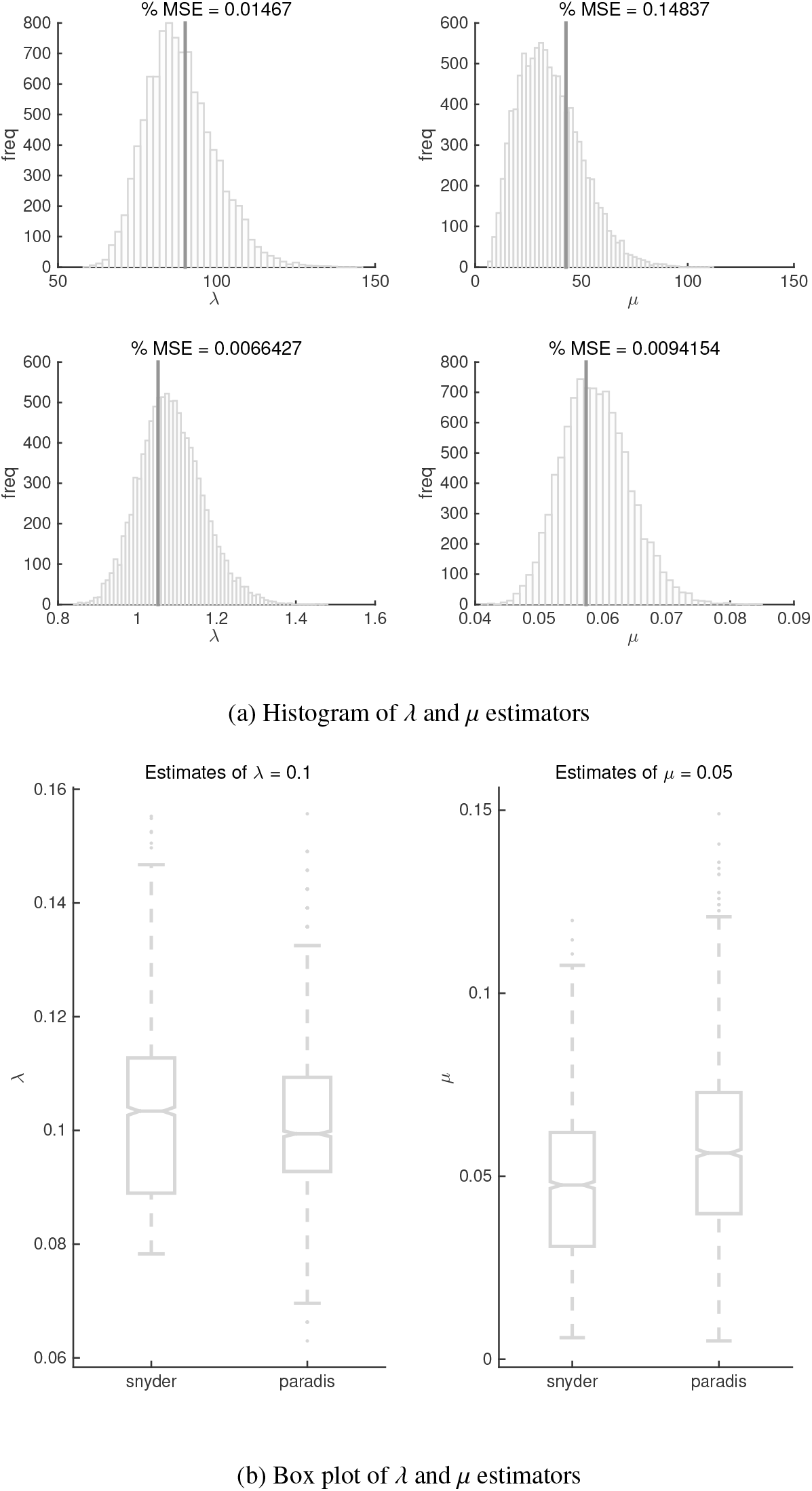
Estimation of the constant rate BDP. The constant rate BDP is simulated and then estimated with the SF. Panel a) is a histogram of these estimates across 10^4^ simulated trees with [*m_i_, n*] = [100, 200]. Simulations are done under the *σ–ρ* parametrisation and converted to *λ–μ* for ease. The top subfigures are for a high death rate setting (priors between [0.01, 0.99] for ρ and [0.01, 100] for *σ*) and the bottom ones for a low setting (priors between [0.01, 0.1] for ρ and [0.1, 2] for *σ*). True values are given in dark grey and the % MSE given as a measure of accuracy. Panel b) is a boxplot comparing the Snyder and Paradis inference methods for the constant rate BDP on 2724 trees with *n* = 200 tips with *m_i_* = 30 grid points selected for both methods. Uniform priors are now directly over [0.01, 0.1] for *λ* and [0.005, 0.5] for *μ* since we want to ensure a fair comparison between methods.

To get an idea of the relative performance of the SF we compared our results with another parametric, least squares method. Developed by Paradis [20], this technique constructs an empirical cumulative distribution function (CDF) and then fits a theoretical, parametric CDF using a sum of squares criteria. The empirical CDF, *F_e_*(*t*), is obtained by normalising the lineage through time plot of the observed birth-death tree. The theoretical CDF, *F*(*t*, {*λ, μ*}), is expressed below for some possible value of (*λ, μ*). We derive this by taking the original equations from [20] and substituting expressions from [2] for constant rate BDPs with *D_s_*:= *λ* – *μe*^(*λ*–*μ*)(*T–s*)^.

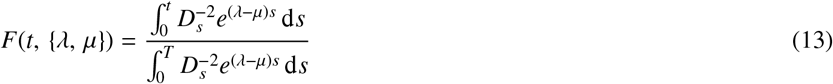

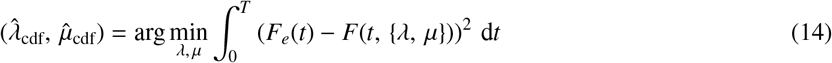

If the search space for (*λ, μ*) is matched to that used in the SF then it is easy to compare the results of the two methods. Since the parameter space is small for the constant rate BDP we were able to find the optimal least squares Paradis estimates 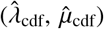 using a brute force search. The integral in equation 13 can be analytically solved. This is given below for completion with the parametrisation *a* = *λ* – *μ*, *b* = *λ*, and *c* = *μe*^(*λ–μ*)*T*^.

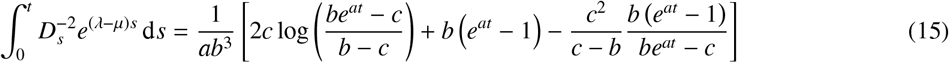

The Snyder-Paradis comparison is given in Figure 1b for 2724 replicate 200 tip rBDP trees. A uniform prior of 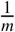 is used across the Snyder grid in the (*λ, μ*) parametrisation to keep comparisons fair. If we used uniform priors on (*σ, ρ*), the actual induced priors for (*λ, μ*) would be non-uniform. The Paradis method achieves a slightly better estimate of *λ* but a noticeably worse one for *μ*. Overall we find the methods very comparable in performance, with the SF being computationally simpler. Both methods are designed to use parametric square error type metrics of performance hence making the comparison sensible.

### 3.2. Snyder Filter BDP Likelihood

We can numerically solve the filter equations from section 2.1 with the rate function from 2.2 to estimate the parameters of BDPs. In section 3.1 we showed this for constant rate BDPs and in later segments of this paper we will extend this to time-varying BDPs. Here we provide and investigate another SF solution. When 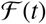 is a standard inhomogeneous Poisson process (so *β*(*t*) is not a function of the lineages), Snyder and Miller showed that the filter could be solved to give the posterior according to [23]:

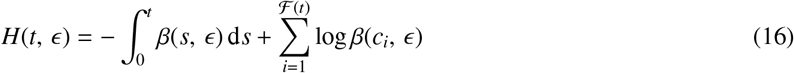

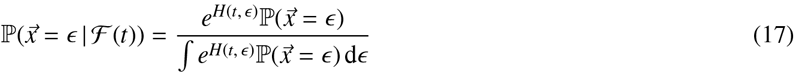

We use *ϵ* for an arbitrary value of the parameter vector 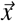 so that enumerating the above expressions across all *ϵ* gives the complete posterior distribution. The *e*^*H*(*t,ϵ*)^ acts as a likelihood function and expression 17 is simply Bayes law. If the *β*(*t*) function admits closed integrals then this solution is completely analytic.

However, our BDP problem is self-exciting in addition to being inhomogeneous. We therefore modify these expressions to account for the extra dependence *β*(*t*) has on the number of existing lineages in the rBDP. If *c_i_* are the observed event times for 1 ≤ *i* ≤ *n* – 1 then the rBDP is inhomogeneous between event times (these are the regions in which the SF of section 2.1 has continuous solutions). We can therefore disaggregate the likelihood into interval sums as 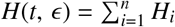 and then use the appropriate self-exciting birth rate *β*(*s, ϵ, i* – 1) for times *s* in this interval starting with *i* – 1 lineages. This gives us the expressions below. In equation 19 we use the posterior from the interval ending with *i* – 1 lineages, 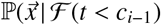, as the prior for the subsequent interval by the Markov property.

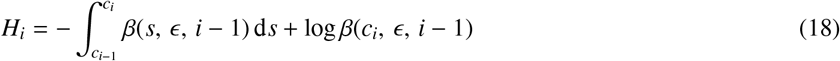

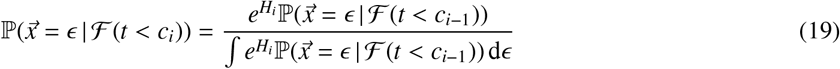

The constant rate BDP allows for closed form solutions to the integral in equation 18 thus making this method useful for this specific inference problem. We obtain the following relation for *H_i_*. The *ϵ* values correspond to different combinations of the parameter set (*σ, ρ*). Our only source of error is the parameter discretisation that implicitly features due to the sequential Markov normalisation. We present a sampled version of this likelihood in Appendix A.

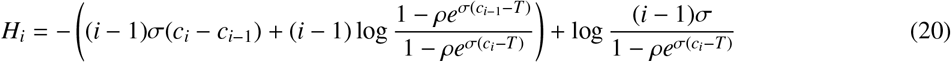

We compared this analytic solution to the standard SF one in equations 1–3. The advantage of the former is that it is not susceptible to errors from numerical integration. The inclusion of explicit exponentiation can lead to minor numerical issues but the Markov decomposition we proposed in 19 resolves this with sequential normalisation. The results of the comparison are given below. We simulated *M* = 10^3^ trees with 100 tips under a high *μ* setting to emphasise any difference between the methods. Figure 2a shows that the distribution of estimated parameters from each method is very similar. As expected, the analytic solution was slightly more accurate, achieving relative % MSE values over [*ρ, σ*] of [10.139, 7.1176] to [11.742, 7.3410] from the original Snyder solution. We found that the correlation between the posteriors calculated by both methods was consistently above 0.988 and that the closeness of the estimates from the methods improved at lower μ settings.

**Figure 2:**
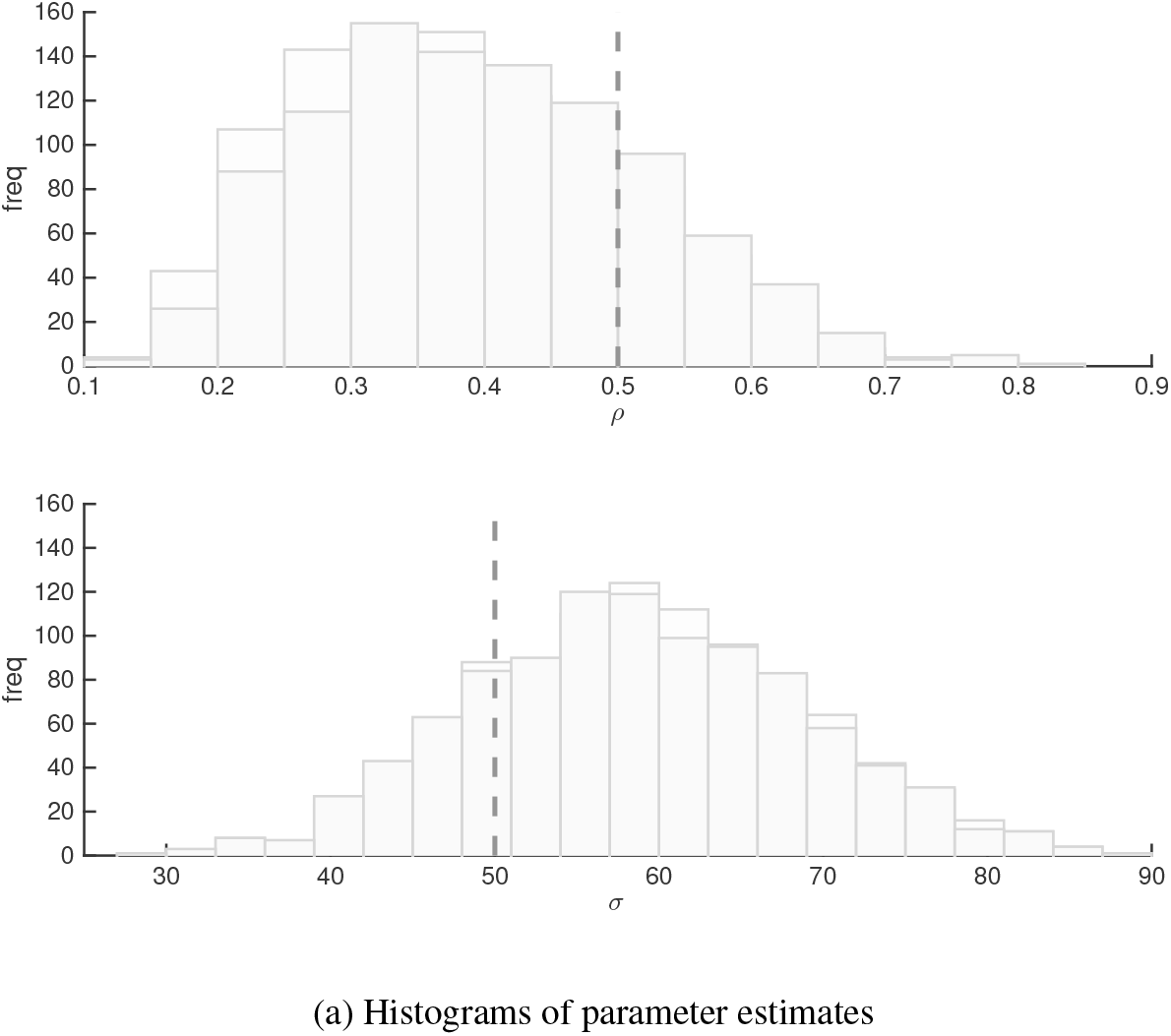
Comparison of constant rate birth-death estimators for the numerical self-exciting Snyder method and the analytic inhomogeneous Snyder solution. The same true parameters are used to generate 1000 trees from which speciation times are obtained. Snyder inference using both the self-exciting and inhomogeneous method is performed using *m_i_* = 100 points within 0.01 – 0.99 for *ρ* and 0.01 – 100 for *σ* (the true *σ* and *ρ* are given as vertical dashed lines). The conditional mean estimates from each method for each tree are then combined into the histograms. The darker grey histogram denotes the analytic method. The posteriors leading to each conditional mean estimate, while not shown, had correlations above 0.988 between the two methods. Simulations were done at *m_i_* = 100, *M* = 1000, *n* = 100 and were representative for other (*σ, ρ*) settings.

Unfortunately the analytic inhomogeneous Snyder solution of equations 16–17 is not useful for time-varying rate BDPs because the integrals are not closed for these problems [2]. In such cases there is no advantage in using this method over the original Snyder solution. Additionally, this inhomogeneous solution is not as flexible to more complex dynamics. Consequently, we only employ the original SF method for solving inference problems in the rest of this text. However, we do make use of the likelihood expressions derived from this method for later comparisons on both constant and time-varying rate BDPs.

In particular, we now use the log likelihood from equation 20 to determine what conditioning assumptions the SF is making for BDPs. As noted by Stadler [30] there are seven different BDP likelihoods even for the simplest constant rate BDP. These arise from conditioning the BDP on different quantities and can affect the accuracy and comparability of estimates obtained from different BDP inference schemes. We examine the likelihood that is implicitly solved by our Snyder inference procedure and compare it to the seven outlined by Stadler.

We start by examining a fine grid over the (*σ, ρ*) parametrisation of the constant rate BDP with *m*_1_ = *m*_2_ = *c* = 150 and generate a rBDP tree with parameters at the median of this grid. We evaluate the log likelihood functions at each point on this grid over the branching times of this observed tree and then marginalise for each parameter. We plot all seven of the marginal log likelihood sets outlined by Stadler [30] in Figure 3a below. To be consistent with the definitions in Stadler, we converted our rBDP speciation times so that they are referenced backward in time and appropriately adjusted our initial conditions to account for when a likelihood function assumed a process starting at the crown instead of the tree root. Figure 3a clarifies that conditioning can make a noticeable impact on the likelihood.

**Figure 3:**
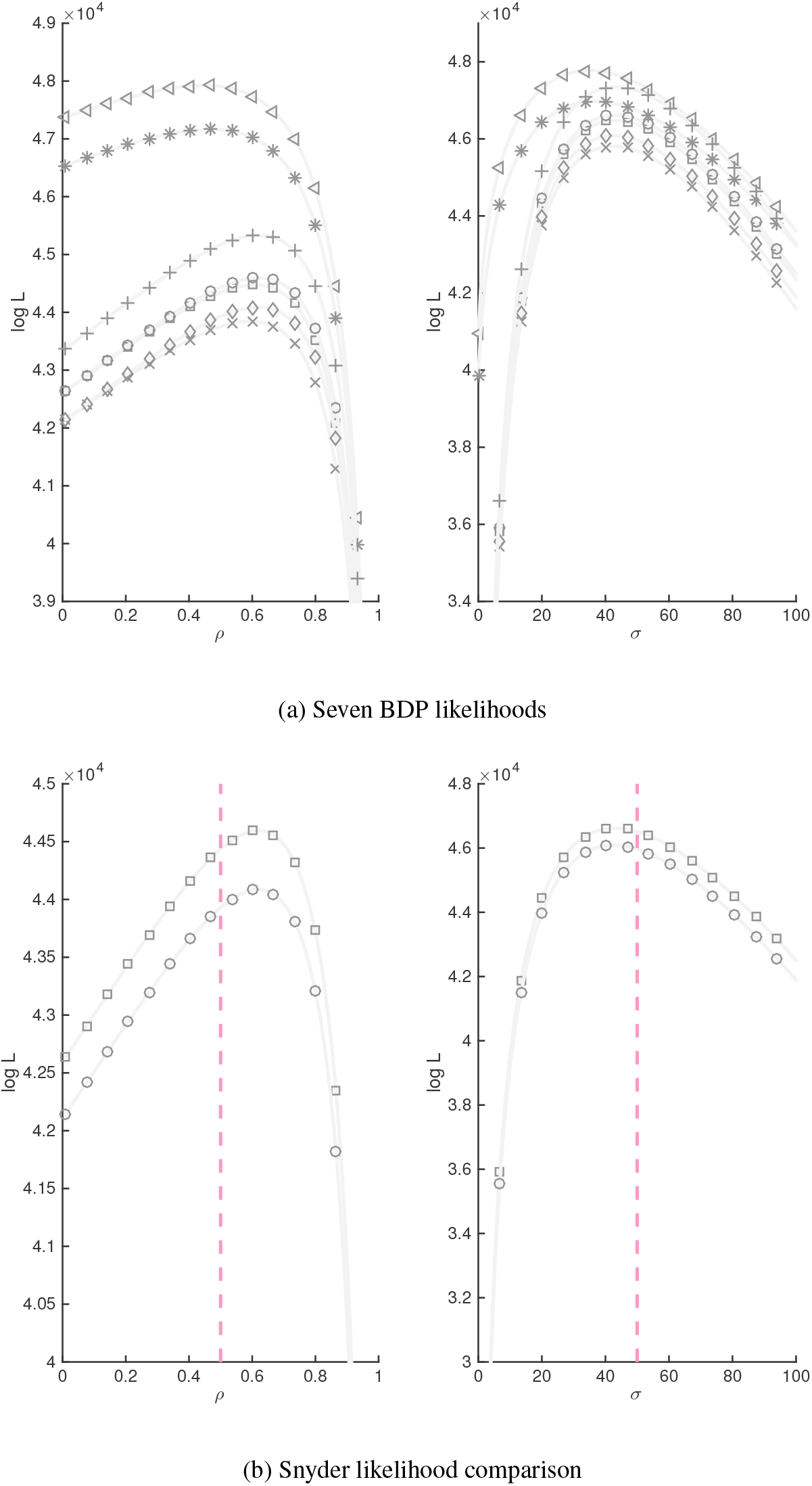
Comparison of constant rate birth-death likelihood functions from Stadler and Snyder. The seven Stadler likelihoods and the Snyder likelihood from the inhomogeneous method are examined over a grid with *m*_1_ = 150 points within 0.01 – 0.99 for *ρ* and *m*_2_ = 150 points within 0.01 – 100 for *σ* (the true *σ* and *ρ* are at the range medians). Panel a) shows the Stadler log marginal likelihoods for each parameter. Consistent markers indicate marginals from the same joint likelihood. All the curves are based on the same tree data, adjusted as necessary to meet various conditions. Panel b) compares the Snyder analytic inhomogeneous marginal log likelihoods to the 2nd and 5th Stadler ones. The light grey curves are the Stadler likelihoods. The squares are the Snyder likelihoods for conditioning on the survival to *T* of the root lineage. The circles are for conditioning on the survival of both crown lineages to *T*. The grey dashed lines are the true parameter values.

We compared the full set of Stadler likelihoods to the analytic SF likelihood on the same tree. Again the relevant adjustments are made to account for time scale and tree conditions. When the SF is applied to a tree that starts with the root, it exactly matches the 2nd Stadler log marginal likelihood provided the correction constant *C* = −*c* log((*n* − 1)!) is added. This correction is equivalent to multiplying the Stadler joint likelihoods with (*n* − 1)! (when marginalising we sum the log of this factor a number of times equal to the number of nuisance grid points – hence the *c* factor). As noted in [30] this means the SF is computing a likelihood on branching times because for a given (observed) branching time vector there are (*n* − 1)! different (equally likely) oriented trees.

The 2nd Stadler likelihood is for a BDP conditioned on just survival to *T* (the present). It makes sense that the SF computes this as the Markov birth process rate *β*(*t*) depends on *P*(*t, T*), a function that encodes survival time. If tree data starting from the crown or MRCA is applied then interestingly the SF matches the 5th Stadler likelihood with the same correction constant *C*. The 5th likelihood is for a BDP conditioned to start at the crown with both initial lineages surviving until *T*. The match of the marginal log likelihoods between these Stadler functions and the Snyder one is given in Figure 3b. The Stadler values are in light grey and the corrected Snyder ones given as square and circle markers for the 2nd and 5th likelihoods respectively.

The equivalence shown here is important for three reasons. Firstly, it confirms our Snyder implementation. Secondly, it makes comparison of results from our method and others in the literature more transparent and valid. For example, later in this manuscript we compare the Snyder estimates to those from an MCMC method by Hohna *et al* [8] [31] (which makes use of the Stadler likelihoods but in a time-varying sense). Since we know what the SF is conditioning on we can ensure we use the relevant Hohna *et al* conditions. Keeping the data and likelihood functions consistent between methods means that any difference in inference results will directly reflect differences only in the techniques themselves. Lastly, Stadler [30] noted that different likelihoods can lead to different estimate accuracies and recommended using the 2nd and 5th likelihoods due to their robustness and accuracy. The fact that these are exactly the likelihoods solved by the SF is quite promising for our point process approach.

### 3.3. The Yule-Furry Model and the Kingman Coalescent

BDPs and coalescent models offer popular, alternate descriptions of the diversity patterns within a population. Understanding the similarities and differences between BDPs and coalescent process descriptions is important for choosing the appropriate null model for a given dataset [15] [16]. Thus, while we are primarily concerned with BDPs, we make some comments on and comparisons with the fundamental Kingman coalescent [14], which models the diversity in a constant population. Previous comparisons have usually been from the perspective of density functions or inter-event intervals. Stadler [32] [5] showed that a constant rate BDP is only analogous to the Kingman coalescent under the specific sampling condition of 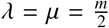, for a tree downsampled from *m* to *n* taxa. Hey [33] found that the true birth-death equivalent to the Kingman coalescent (provided a Volz rate correction is applied [34]) is achieved if birth and death events are synchronised. This leads to a Moran type model, also with *λ* = *μ*. Our BDP-coalescent comparisons are from the perspective of the resulting parametric inference problem.

Unlike the BDP, the coalescent is defined in backward time and assumes a population size that is large in comparison to the sample size *n* [35]. We first rewrite the Kingman coalescent equations in forward time to match the BDP and use 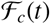 as its observed process that counts the number of tree lineages at *t*. The Kingman coalescent with population *N*(0) then has observable inter-event intervals exponentially distributed with rate 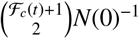. Given 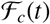 then this interval derives from a homogeneous Poisson process. Consequently, it admits a simple and well-known analytic solution for the estimator of *N*(0). We presented the analytic SF solution for this problem in [18]. The constant rate BDP with *λ* = *μ* however does not lead to a similar homogeneous Poisson process description. This is because the observable rBDP is not homogeneous given 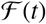, and therefore presents a more complex inference problem. This can be seen by applying L’Hopitals rule to expression 10 but in the original (*λ, μ*) parametrisation.

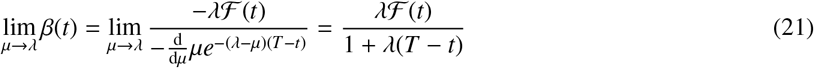

This results as the birth and death events are completely independent, which is in contrast to the Hey model [33]. It also emerges because when conditioned on survival (which is the case for an rBDP to be observed) then the expected number of lineages is not constant despite the equality in the birth and death rates [9]. The impact of conditioning becomes clear if we use the expression given by Hohna [9]. Hohna finds, for all BDPs with *λ*(*t*) = *μ*(*t*), that *P*(*t, T*) is the same as the inverse of the expected number of lineages at *T* conditioned on survival of the true process (hence we use *l*(*t*)). Using Hohna’s expressions in our *β*(*t*) calculation gives the following.

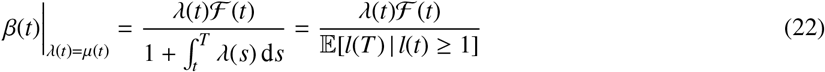

However, the inference problem for the Kingman coalescent can be made identical to that for a BDP if the death rate is set to 0. The resulting model is called a Yule-Furry process [36]. Estimation on this process is with the true tree as the rBDP and BDP are equivalent at *μ* = 0. This allows us to analytically solve the SF equations in a manner that mirrors the solution that we presented in [18] for the Kingman coalescent. Additionally, also as in [18], we will describe the solution robustness to prior specification.

The Yule-Furry process has inter-speciation intervals that are independent and exponentially distributed with rate *iλ* over the period *c*_*i*+1_ – *c_i_* where 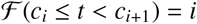. The known maximum likelihood estimate of *λ*, 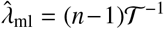 [37]. 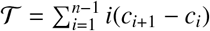 is the sufficient statistic for the Yule-Furry process. If the observed intervals {*c*_*i*+1_ – *c_i_*} for all *i* ∈ {1, *n* – 1} are exponentially scaled by the number of lineages to {(*c*_*i*+1_ – *c_i_*)*i*} = {*δ_i_*} and a new observed process defined 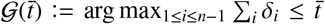, then 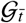 is homogeneous and Poisson. For a homogeneous process Snyder showed that the filter equations can be solved analytically to give the formula 23 below [17]. As usual *ϵ* indicates some arbitrary value of the parameter of interest. This solution is only applicable to the Yule-Furry because of the transformation from 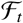 to 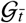. Note that 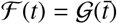 and the conditional mean 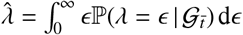.

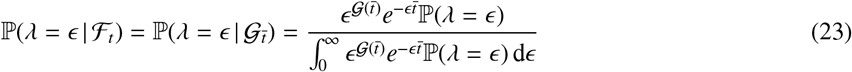

Here 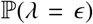 is the prior and the exponentially scaled observed stream ends at 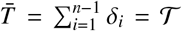. We can obtain the Bayesian sensitivity of the Snyder Yule-Furry solution to prior specification by describing the dynamics of the maximum a posteriori (MAP) estimate of *λ*. The MAP estimate is the mode of the posterior 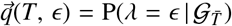. We simplify and solve Snyder’s MAP estimate differential equation: 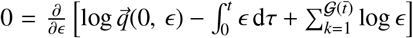 [17] for the Yule-Ferry process to get expression 24, with 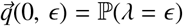.

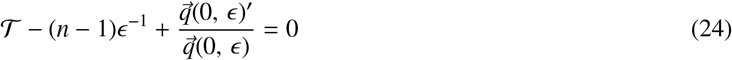

Solving for *ϵ* gives the MAP estimate 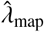. It clearly depends on the logarithmic derivative of the prior, 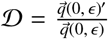. 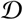 represents the sensitivity of the Bayesian MAP estimate to the prior. For a uniform prior 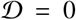 and 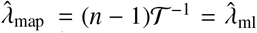. If the prior is a negative exponential: 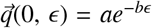 then 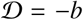 and 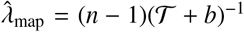. Most importantly, as *n* (and hence 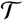) gets large, then for any prior, 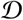, becomes negligible and every 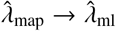. In Appendix A we present a new sampled condition under which a constant rate BDP with *λ* ≠ *μ* is analogous to this Yule-Furry one and thus can admits all the above results.

For completion, we write the analytic Snyder likelihood for the entire observed Yule-Furry process *e*^*H*(*T,λ*)^, as equation 25 below, by directly solving equation 16. Note that the rBDP rate is now parametrised only in terms of *λ*.

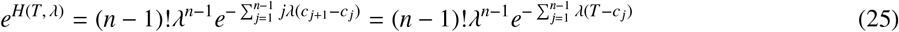

Expression 25 is equivalent to (*n* – 1)!*L*(*λ*) where *L*(*λ*) is the tree likelihood for an rBDP conditioned on *T* with *μ* = 0. This is one of the seven rBDP tree likelihood functions presented in [30] and further confirms the Snyder method.

### 3.4. Time Varying Rate Birth-Death Estimation

We now generalise to estimating BDPs with deterministically time-varying (but not lineage dependent) per lineage birth and death rates *λ*(*t*) and *μ*(*t*). These types of models, while not as tractable as the constant rate BDP, are important for describing complex diversification dynamics such as adaptive radiations or mass extinctions [20]. As mentioned in section 2.2 we assume that the birth and death rates can be parametrised as 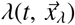 and 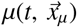 with 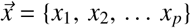 for a *p* ≥ 1 parameter BDP. The vectors 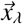 and 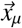 are spanning subsets of the parameter set. This means that both the birth and death rate may be functions of all the *p* parameters or just a subset but each parameter must appear at least once (otherwise it cannot be inferred). Often we will just write *λ*(*t*) and *μ*(*t*) for convenience.

Time varying BDPs admit no explicit inference solutions [20] and so we will numerically integrate the Snyder equations 1–3 across the observed reconstructed process or rBDP. We previously noted that this rBDP is equivalent to a Markov birth process, 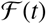, with rate at time *t* ≤ *T*, given in equation 8 and with 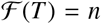. This rate is explicitly written for convenience in equation 26 to show its parametric form and that it depends on nested integrals. This makes the inference problem much more complex than the constant rate BDP one. The rate is still Markov because it only depends on the current count of the rBDP [23].

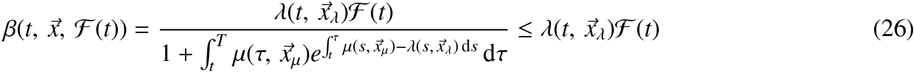

The inference problem is to find conditional mean estimates of the parameters *x_i_*, 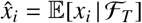. The SF will directly generate the joint posterior 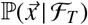 and the estimate 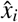 will be obtained by marginalising for this parameter and then integrating the marginalised posterior across its domain.

We investigate two time-varying rate BDP models. The first has a constant death rate *μ*(*t*) = *x*_3_ and an exponentially decreasing birth rate *λ*(*t*) = *x*_3_ + *x*_1_*e*^−*x*_2_*t*^. We call this the Hohna model since he introduced it in [38] to describe a speciation rate which initially starts above the extinction rate and then decays to a critical *λ*(*t*) = *μ*(*t*) state. He used this and various nested simplifications for a model selection problem on empirical ant and snake phylogenies. The second model uses a logistic function for both the birth and death rate so that: *λ*(*t*) = (1 + *e*^−*x*_1_*t*+*x*_2_^)^−1^ and *μ*(*t*) = (1 + *e*^−*x*_3_*t*+*x*_4_^)^−1^. This model, which we refer to as the logistic, was used by Paradis [20] to capture the monotonically increasing or decreasing birth and death dynamics commonly found in the macroevolution literature.

We simulated reference (observed) time-varying rBDPs from each model using the algorithms of Hohna [31]. These are available in the R package TESS [21] and involved inversely sampling the *i*^th^ rBDP speciation time, *c_i_*, from the theoretical CDF of equation 27. *P*_1_(*t, T*) = *P*(*t, T*)^2^*e*^*α*(*t,T*)^ is the probability that a lineage at *t* leaves exactly 1 surviving descendant at *T*. Note that *P*(*t, T*) is the analogous probability for when at least 1 descendant survives to *T*). Here each 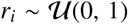 and the tree generated is for *n* lineages at time *T* so that *F*(*T*) = *n*. Where possible, we simulated under true parameter values that matched those found in the previous Hohna and Paradis works.

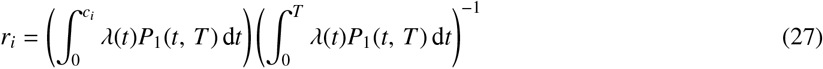

Having simulated the observed data for each time-varying BDP model we then applied the SF and compared its estimates to the true value. To get an idea of relative performance of our method, we also analysed the simulated trees with a recent fixed tree adaptive MCMC inference method. Developed by Hohna *et al* [31] [21], and also included in the TESS package, we used this method as a modern benchmark for time-varying BDP estimation. This Hohna technique also accommodates incomplete sampling and density dependence.

Since the root of a tree is less likely to be observed in real life, we conditioned our rBDP trees to start from the crown or most recent common ancestor. Our starting time is therefore *c*_1_ instead of 0 and 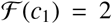. For these inference problems, the Hohna MCMC method samples from the likelihood function given in equation 28. This is the time-varying form of the 5th Stadler likelihood mentioned in section 3.2 for constant rate BDPs.

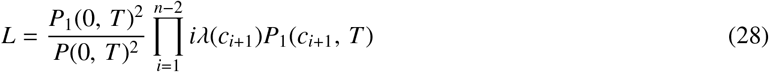

For all models we simulated *M* = 100 replicate rBDP trees with *n* = 100 tips. We estimated model parameters from the simulated trees using both the Snyder and Hohna techniques. To keep comparisons fair we used the uniform priors across the same parameter ranges for both methods. For the SF we used probability 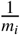 for the *i*^th^ parameter grid, with *m_i_* = 20 for the Hohna model and *m_i_* = 15 for the logistic model. We checked the convergence of the MCMC method by using the Geweke statistic [39] and by examining the autocorrelation function of the MCMC samples.

Figures 4a and 5a below show the resulting combined marginal posteriors from both methods. These are obtained by combining and smoothing histograms of samples generated from the marginal posteriors estimated from each tree. While there is some bias in the estimates of certain parameters, it is clear that all the true values are within the cover of the posteriors. This bias emerges because time-varying BDPs present a difficult inference problem [20]. Most importantly both methods lead to well matched marginals, validating the performance of the SF. We confirm this further by plotting the Hohna and Snyder marginal likelihoods across each *m_i_* point parameter grid. These are given in Figures 4b and 5b. Within numerical tolerances the likelihoods from both methods are identical. For illustration, we converted the Snyder posteriors from each tree into an estimated birth and death rate trajectory. We present a summary of these trajectories in Figures 6a and 6b. The Hohna death rate is accurately measured but the birth rate is initially underestimated. In contrast, the logistic birth rate is well covered but its death rate is initially overestimated. Note that estimates of both the parameters and the rates will improve as *n* increases.

**Figure 4:**
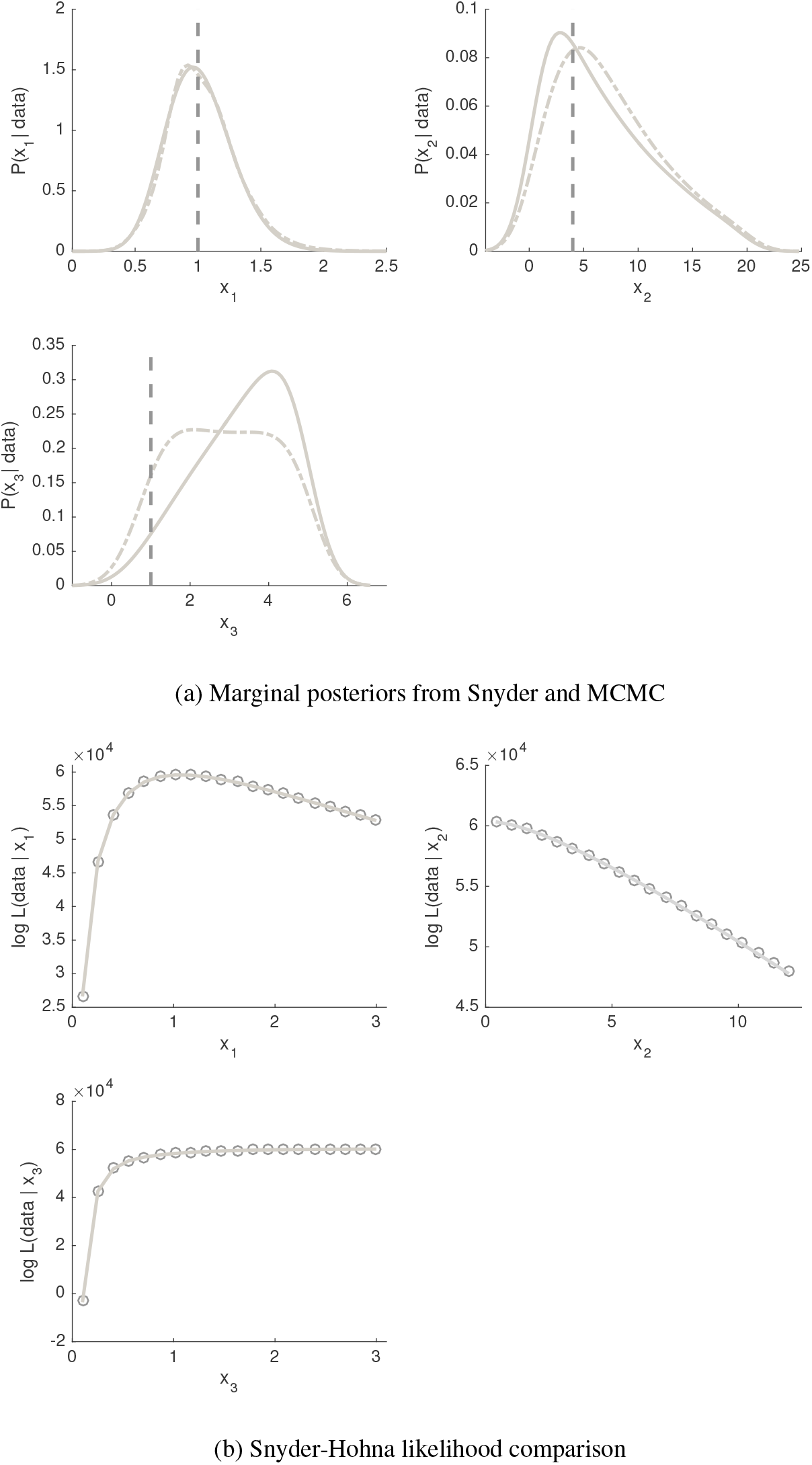
Snyder and MCMC estimates for the Hohna time-varying BDP model. We simulated *M* = 100 trees with *n* = 100 tips and then estimated the underlying parameters of the model using the SF and Hohna MCMC methods. We used uniform priors spanning [0.1, 0.4, 0.1] to [5, 20, 5] for the 3 parameters respectively. We ran the Snyder method on a grid with [*m_i_, m*] = [20, 20^3^] points. Panel a) shows the smoothed marginal posteriors obtained by aggregating samples from all the tree posteriors and then applying a normal smoothing kernel. The MCMC posteriors are in solid grey and the Snyder ones in dashed grey. The dark vertical dashed line is the true parameter value. Panel b) compares the Snyder and Hohna MCMC marginal likelihoods. We evaluated the relevant joint likelihood functions for both methods on a given tree over the 20^3^ grid points, and then marginalised. The solid grey curves are the Hohna likelihoods while the circle markers are the Snyder ones.

**Figure 5:**
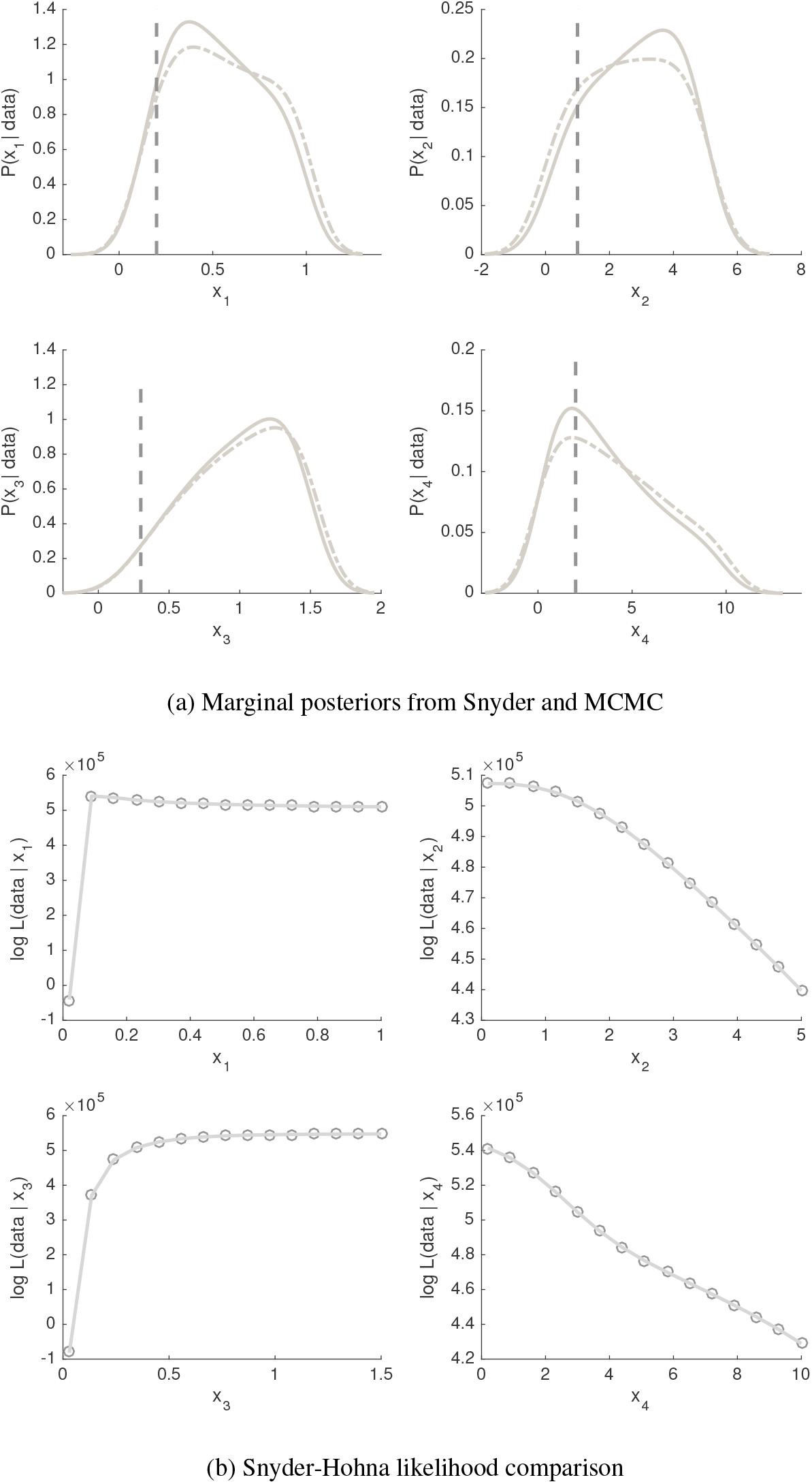
Snyder and MCMC estimates for the logistic time-varying BDP model. We simulated *M* = 100 trees with *n* = 100 tips and then estimated the underlying parameters of the model using the SF and Hohna MCMC methods. We used uniform priors spanning [0.02, 0.1, 0.03, 0.2] to [1, 5, 1.5, 10] for the 4 parameters respectively. We ran the Snyder method on a grid with [*m_i_, m*] = [15, 15^4^] points. Panel a) shows the smoothed marginal posteriors obtained by aggregating samples from all the tree posteriors and then applying a normal smoothing kernel. The MCMC posteriors are in solid grey and the Snyder ones in dashed grey. The dark vertical dashed line is the true parameter value. Panel b) compares the Snyder and Hohna MCMC marginal likelihoods. We evaluated the relevant joint likelihood functions for both methods on a given tree over the 15^4^ grid points, and then marginalised. The solid grey curves are the Hohna likelihoods while the circle markers are the Snyder ones.

**Figure 6:**
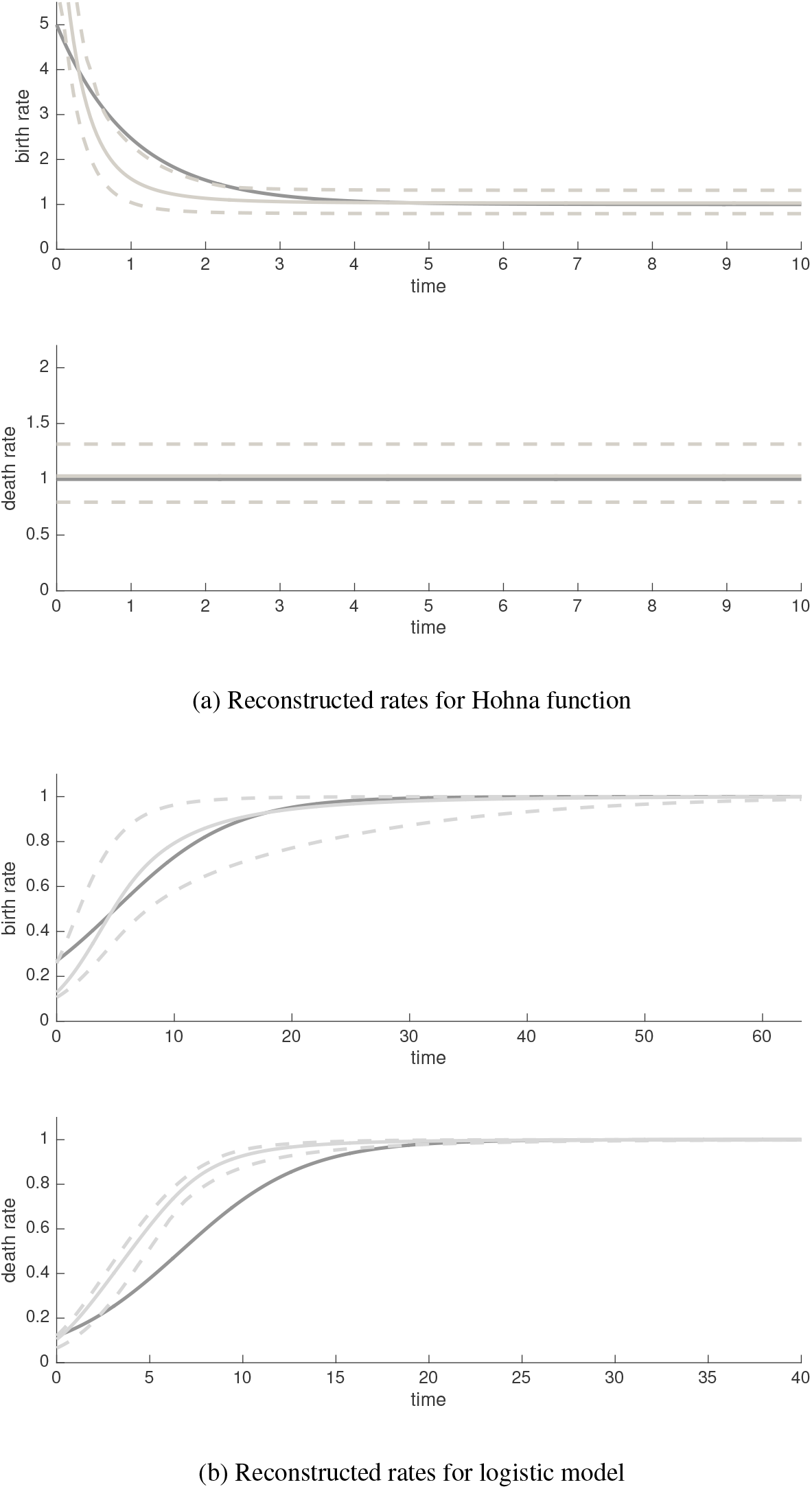
Reconstructed birth and death rates for time-varying BDPs. The SF is applied to *M* = 100 replicate *n* = 100 tip trees simulated from the Hohna and logistic models. The joint posterior from each tree is used to obtain a reconstructed birth (*λ*(*t*)) and death rate *μ*(*t*). Panel a) provides a summary of these estimated birth and death trajectories for the Hohna model whilst panel b) illustrates the logistic model trajectories. The SF settings match those in Figures 4a and 5a respectively. In both panels the solid grey line is the mean of the estimated trajectories while the dashed lines enclose 95% of all trajectories. The dark solid line is the true rate.

### 3.5. Birth-death Estimation with Empirical Data

We extend our analysis of time-varying BDPs to empirical data. As the SF is flexible and easily implemented it should be useful for model selection problems. We test that assertion on the Australian Agamid lizard dataset from Harmon *et al* [40], which is known to be almost completely sampled (above 93%) at the species level [22]. Previous work by Rabosky and Lovette (herein referred to as simply Rabosky for convenience) [22] analysed 4 (nested) birth-death models and found that the dataset was best described by a constant death rate, *μ*(*t*) = *x*_3_ and an exponentially decreasing birth rate, *λ*(*t*) = *x*_1_*e*^−*x*_2_*t*^. This supported a model for declining diversification. We apply the SF to the same nested parametric models used by Rabosky, and use the same time scaled Agamid tree as our observed process. These models, with parameters *x_i_* are: i) constant birth-death, *const: λ*(*t*) = *x*_1_, *μ*(*t*) = *x*_2_; ii) time-varying speciation, *spvar: λ*(*t*) = *x*_1_*e*^−*x*_2_*t*^, *μ*(*t*) = *x*_2_; ii) time-varying extinction, *exvar: λ*(*t*) = *x*_1_, *μ*(*t*) = *x*_3_(1 – *e*^−*x*_2_*t*^) and iv) time-varying birth-death, *bothvar: λ*(*t*) = *x*_1_*e*^−*x*_2_*t*^, *μ*(*t*) = *x*_4_(1 – *e*^−*x*_3_*t*^).

Since there are no true parameter values for this problem we use the maximum likelihood estimates (MLEs) from Rabosky’s method as our reference. We recalculated these using the R package LASER [41]. LASER uses the function ‘optim’ with the ‘L-BFGS-U’ method to maximise a likelihood function from Nee *et al* [2]. In order to protect against local maxima the optimisations were repeated 100 times. We perform model selection with estimates from both the Rabosky and Snyder methods by using a least squares technique developed by Paradis [20]. This method involves converting the tree data into an empirical cumulative distribution function (CDF), *F_e_*(*t*) (essentially a scaled version of 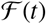. A theoretical CDF, *F*(*t, ϵ*) for some arbitrary value of the parameter vector, 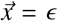, is also defined as in equation 29. The accuracy of a model is assessed by the square error: 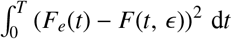.

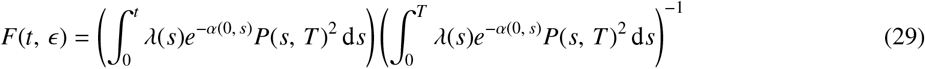

We used this Paradis metric since it was simple, easy to visualise and fits directly to the empirical data of interest. More complex model selection techniques were found unnecessary for this problem. Additionally, sometimes using such techniques, like the Akaike information criterion, can be misleading for BDP problems [42] [38]. We calculated the Paradis metric across the 4 models and scaled the resulting values by the maximum model value (the result is a value between 0 and 1). Our model selection results are given in Figure 7a. The main plot compares the Snyder based 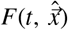 for each model (light) against the empirical *F_e_*(*t*) (dark). The SF was run for the Agamid *n* = 69 tree with uniform priors of 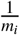 on each parameter for every model. The grid sizes ([*m_i_, m*]) were: [30, 30^2^], [30, 30^3^], [30, 30^3^] and [20, 20^4^] for *const, spvar, exvar* and *bothvar* respectively.

**Figure 7:**
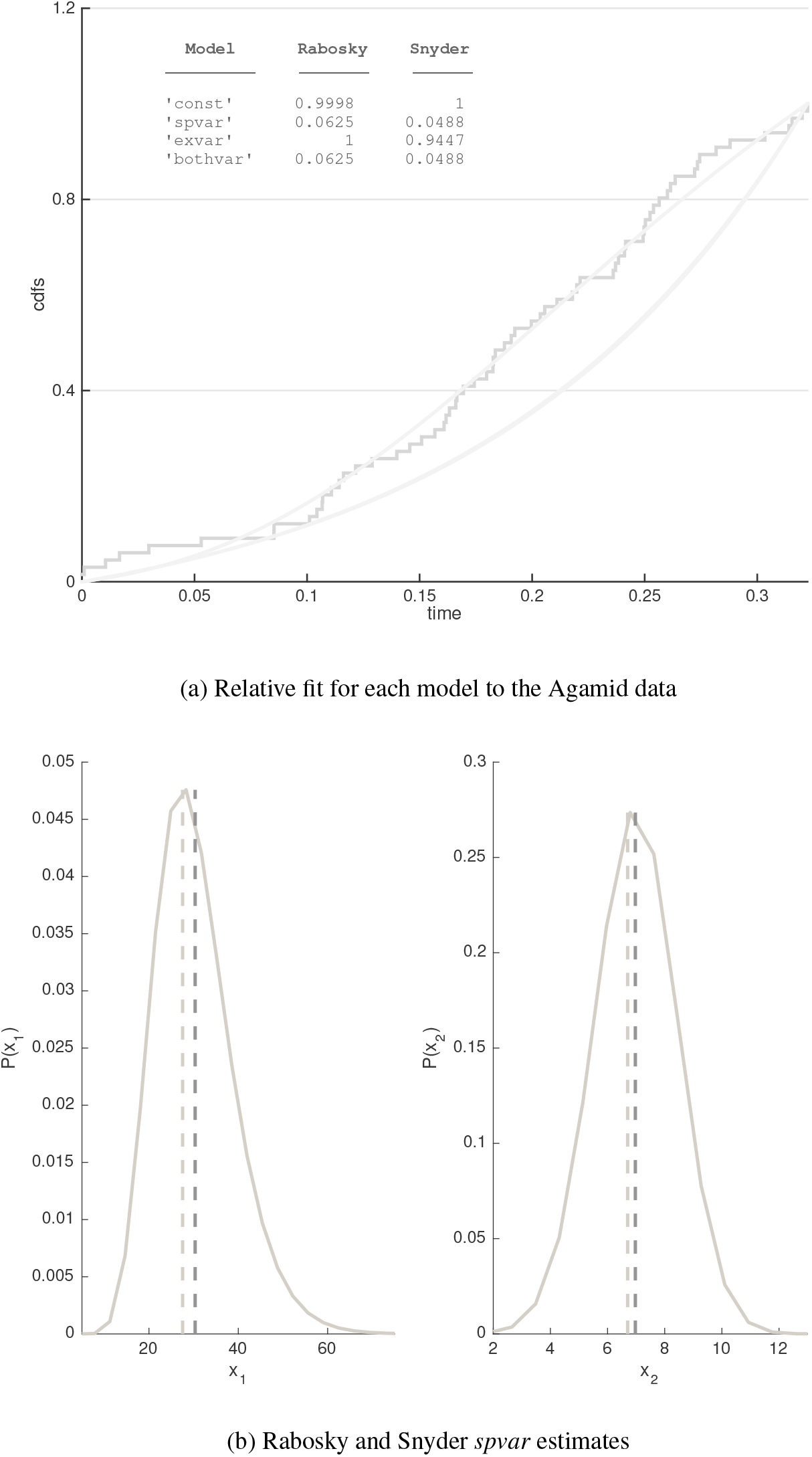
Comparison of several time-varying BDP models on the Agamid dataset. The SF and Rabosky method were applied with the *const, spvar exvar* and *bothvar* models to the *n* = 69 tip Agamid tree. All models were examined with *m_i_* = 30 except for *bothvar* which had *m_i_* = 20. Snyder priors were uniform at 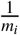 over the ranges [0 – 10, 0 – 1], [1 – 100, 1 – 25, 10^−3^ – 0.01], [0.01 – 10, 0.01 – 10, 10^−3^ – 0.01] and [1 – 100, 1 – 25, 0.01 – 1, 10^−3^ – 0.01]. Panel a) gives the Snyder CDFs (light curves) against the empirical CDF, *F_e_*(*t*) (dark stepwise function). The *const* and *exvar* fit poorly (lie under *F_e_*(*t*)) while the *spvar* and *bothvar* show a much better and identical fit (the curves are overlaid). The inset table gives the relative fit to *F_e_*(*t*) based on the relative ratio of the Paradis metric. Results support *spvar* as the best model. Panel b) therefore examines the Snyder *spvar* posteriors (solid light curve) against the Rabosky MLEs (dark dashed line). The light dashed line is the Snyder conditional mean estimate. No comparison is provided for *x*_3_ since it seems irrelevant (its posterior matches its prior).

The *spvar* and *bothvar* are the top grey curves, and seem to fit the empirical data equally well. Since the *spvar* has less parameters, it is the best model for the Agamid data. The other models are below the empirical CDF and seem to fit poorly. The inset table lists the scaled Paradis metric for both the Rabosky and Snyder inference. Observe that the relative fit given to each model is essentially consistent across the methods and matches that in the original Rabosky analysis [22]. By confirming the *spvar* model description, the SF is also supporting a declining diversification rate hypothesis for the Agamids. Figure 7b examines the Snyder marginal posteriors for the best fit *spvar* model. The dark dashed line indicates the Rabosky MLE from LASER (we were not able to get confidence intervals for the Rabosky estimates). Note that the Snyder posteriors are strongly matched to the Rabosky estimates. The posterior for the death parameter, *x*_3_, is not shown because it remained unchanged from the set prior. This suggests that either the death rate is not an important explanatory variable or that the Agamid rBDP may not be sufficiently informative to allow death rate dynamics to be inferred. The essentially zero death rate was also found in the original Rabosky work on this dataset [22]. The SF therefore provides evidence for a more concise model than *spvar* that excludes death events.

## 4. Discussion

In this paper we have introduced the SF as an alternate approach to BDP rate estimation problems. We have shown that if we exploit the equivalent Markov birth process statistics of a BDP’s associated rBDP, then the SF can be used to compute the posterior distribution of its parameters. This method only relies on having a parametric description of the birth and death rates and a means of calculating the probability that a lineage alive at time *t* leaves at least one descendant at *T*, (*P*(*t, T*)). Given such minimal requirements, the SF provides a flexible, useful and accurate inference scheme. We subsequently summarise the evidence of this, which was provided throughout the main text.

We started by using the SF to estimate the parameters of a constant rate BDP. We found that its estimates were accurate and improved with decreasing death rate. These trends matched the literature. We found the SF performance to be comparable to a least squares optimisation method by Paradis [20], without requiring a search for minima itself. One benefit of the SF is its lack of susceptibility to local optima. We also used Snyder theory to derive (for the first time, to our knowledge) the analytic MMSE estimator for the Yule-Ferry process. This was analogous to the Kingman coalescent solution that we have previously presented in [18]. Interestingly, in Appendix A we found that under a new and specific homochronous sampling condition the constant rate BDP behaves like a Yule-Furry process with parameter *λ* – *μ* and so also admits an equivalent analytic estimator.

We then observed that by conditioning on the number of lineages between the branching events of the rBDP we could adapt an analytic SF solution, known to apply to inhomogeneous Poisson processes. We easily obtained marginally more precise results than the basic SF implementation for constant rate BDP estimation, using this method. Unfortunately, this more analytic technique is not as usable for more complex BDPs. However, this method revealed a likelihood function that the SF was implicitly solving. Stadler [30] stated that seven distinct BDP likelihoods exist depending on what rBDP characteristics are used for conditioning. She examined all of these for the constant rate BDP and found that the 2nd and 5th likelihoods were the most robust and hence useful for study. We examined the implicit Snyder likelihood and found that it was solving the 2nd and 5th Stadler likelihoods, a property likely inherited from the Nee *et al* function *P*(*t, T*) [2]. This confirmed the SF as a useful and valid BDP inference technique.

This likelihood robustness refers to a mismatch which can often occur in BDP analyses. If one simulates a tree with a fixed size *n* then the correct likelihood function is the 7th. This conditions on *n* and places a uniform prior on tree stem age. However, conditioning on survival is important to get more accurate estimates, especially at higher extinction rates. This creates an expected issue, as it is not advised to condition on both survival and *n* [30]. Luckily, Stadler found that the 2nd and 5th likelihood, which condition only on survival at either the root or crown, leads to as accurate estimates as the 7th despite the data being simulated under the *n* condition. This implicitly features in this work since simulations are often done under the *n* condition in order to fix the information content of the tree (more branching events leads to sharper estimates). This allows a better evaluation of an inference method than for example simulating to survival at a fixed *T*. Since the SF uses the robust likelihood functions, then there is no issue. This approach to testing a survival based inference method on *n* conditioned data is not uncommon [43].

We then extended our SF analysis to time-varying BDPs. We simulated two models from the literature: one introduced by Hohna [38] that describes a BDP with an exponentially declining speciation and constant exinction rate, and one used by Paradis [20] featuring logistic birth and death rates. We applied the SF to data simulated from both models and compared the results with a modern fixed tree MCMC method developed by Hohna [21]. For both models, both methods gave very similar marginal posteriors and were found to be solving identical likelihood functions. This confirmed the SF performance for time-varying BDPs as well as clarified a key relative advantage of our method. The Hohna method often required multiple runs to avoid poor convergence and could give different results on the same data. The SF does not have convergence issues, and is a deterministic method. Given the data, the SF will always produce the same posterior. In terms of computational speed, we found both methods comparable. More discussion of the Snyder computational and methodological complexity can be found in Parag and Pybus [18].

Having tested the SF on simulated trees we investigated a model selection problem on an empirical phylogeny of Australian Agamids. This dataset was first analysed by Rabosky and Lovette [22] and we used their original tree and compared our results directly with their MLEs. Both methods, under a Paradis CDF least squares metric [20], supported models that featured declining birth rates and constant death rates. The SF marginal posteriors for the selected model were found to match the Rabosky MLEs very well (the Rabosky point estimates were near the mode of the Snyder marginals). Thus the SF performs well on empirical datasets. The Rabosky MLE for the death rate was 0 while the SF produced a posterior that matched the prior. The SF therefore deems this parameter redundant. The suggests the data is not informative enough or that an assumption in the BDPs has been violated. This can likely be resolved with more generalised models or the addition of fossil data [12].

The SF presents a capable alternative BDP inference technique that, within the numerical tolerances of integration and parameter discretisation, provides exact MMSE estimates by directly solving for the joint parameter posterior. It is simple, deterministic and does not suffer from stability issues like local minima or poor convergence. Most importantly it fully exploits the Markov birth nature of rBDPs, which should allow easy generalisation to more complex BDPs. All that is required is a means of calculating *P*(*t, T*). Some obvious extensions that we can mention pertain to genealogical uncertainty, incomplete sampling and density dependence. This work has only focused on the fixed tree problem. To account for tree uncertainty we could likely sample a set of trees describing this uncertainty, run a SF on each and combine the results in a Bayesian manner. This would be similar to combining multiple Poisson streams and is a generalisation made possible by taking the Markov birth process approach [23].

The dependence on *P*(*t, T*) is a key point for incorporating incomplete sampling. We have already shown how to accommodate homochronous sampling (also known as uniform taxon sampling) by replacing *P*(*t, T*) with the probability that a lineage leaves at least 1 descendant at *T* and is sampled *P_ν_*(*t, T*) (see Appendix A). As long as a sampling process admits a description for *P_ν_*(*t, T*) then the SF can be easily extended to these scenarios. We further note that the extension for density dependence is trivial. Density dependence means that the birth and death rates now become functions of the number of lineages. However, the rBDP already involves linear lineage dependence since 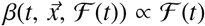. If, in addition, the birth and death rate become functions of 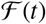 all this means is the structure of *β* becomes more complex. The inference problem, however, is unchanged since 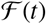 is known (observed data) and the parameter space is still the same. Consequently, as long as the functional dependence can be explicitly written as 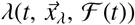 and 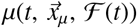 then density dependence is easily accommodated.

Thus we believe that the SF has promise as a BDP inference technique. It is simple and flexible and allows direct computation of the posterior of estimated parameters. Its approach involving equivalent Markov birth processes and ordinary differential equations is fresh to this field and allows a new direction from which estimation insight can be derived. Since it focuses on parametric estimation, we envision the SF as useful for learning about a dataset in a phenomenological manner or comparing different parametric birth-death rate models in an attempt to learn problem physics. Our future work will look to extend this method to handle more difficult problems such as multitype BDPs or BDPs with protracted speciation, with inclusion of genealogical uncertainty.

## Acknowledgments

This work was supported by the European Research Council under the European Commission Seventh Framework Programme (FP7/2007-2013)/European Research Council grant agreement 614725-PATHPHYLODYN.

## Appendix A. The Sampled Birth-Death Process

Throughout the main text we have assumed complete sampling. Whilst the impact of sampling strategy is outside the scope of our work, we provide some analysis of sampled BDPs to show how easily the SF can accommodate such processes. We focus on incomplete homochronous sampling, which includes each extant lineage with probability (or sampling fraction) *ν* < 1. [5]. Since *ν* acts as an additional determinant of what lineages we will see in the sampled rBDP (deaths are the other controlling factor), our rate of producing observed rBDP lineages no longer depends on just *P*(*t, T*) (see equation 6). Nee *et al* [2] noted that the function *P*(*t, T*) simply needs to be replaced with a *ν* sampled version *P_ν_*(*t, T*). For the time-varying BDP this was derived by Morlon *et al* [7] in reverse time. We reproduce their function in forward time using the transformations described in the supporting information of their work. This is given in equation A.1 with 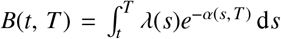. The importance of this expression is the very visible effect of *ν*. When *ν* = 1 this yields the equivalent definition for *P*(*t, T*) given by Kubo and Iwasa [11]. The sampled rBDP rate, *β_ν_*(*t*), can be calculated using equation A.2.

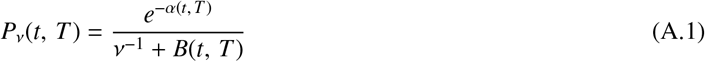

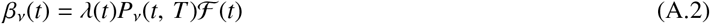

For the constant rate BDP, *λ*(*t*) = *λ* and *μ*(*t*) = *μ* for 0 ≤ *t* ≤ *T*, and equation A.1 reduces to the well known expression found in [2]: *P_ν_*(*t, T*) = *ν*(*λ* – *μ*) (*νλ* + (*λ*(1 – *ν*) – *μ*)*e*^−(*λ–μ*)(*T–t*)^)^−1^. Combining the above expressions with those from the completely sampled case gives the inequalities in A.3. The second half of this relation comes from observing that 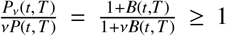. The sampled rBDP therefore has a rate that is between that of a completely sampled rBDP and an rBDP with scaled birth rate of *λ*(*t*)*ν*^−1^. The latter is equivalent to a Bernoulli thinning of a completely sampled rBDP. Additionally, if *ν* is very small, we observe that 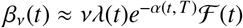 and so the rate of observable informative events is low and directly proportional to the sample probability.

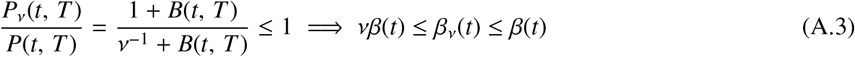

The sampled rBDP rate, *β_ν_*(*t*), is still that of a self-exciting Markov process and so the usual Snyder equations apply to the incompletely homochronously sampled time-varying BDP. The inference problem given *ν* is essentially the same. In fact in the constant rate case, we can assume complete sampling and run the Snyder equations as usual to estimate parameters *x*_1_ = *λ* and *x*_2_ = *μ* as 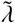 and 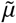. We can then obtain the true MMSE estimates which account for incomplete sampling using v by the transformation given by Stadler [5]: 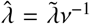 and 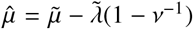. Note that if *ν* is not known then it cannot be estimated simultaneously with 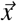 from 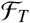 and more information is required [5].

Since the homochronously sampled BDP fits within the Snyder inference framework, a likelihood function similar to those derived in section 3.2 can be written. We simply replace *β*(*t*) with the equivalent *β_ν_*(*t*). We do this for the constant rate BDP and achieve an altered form of equation 20 given below.

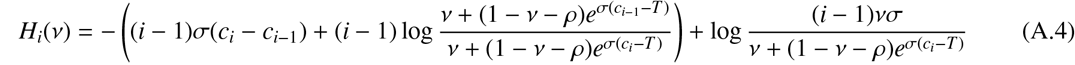

To complete our Snyder analysis of homochronous sampling, we mention an interesting case of the sampled constant rate BDP that admits a solution similar to the Yule-Furry and Kingman coalescent ones from section 3.3. Stadler [32] showed that the sampling condition 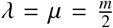 can be used to make a sampled constant rate BDP behave like a Kingman coalescent. Our condition is, however, different and we do not require equal birth and death rates. We sample such that *ν* = 1 – *ρ* in the (*ρ, σ*) parametrisation of the constant rate BDP. In this case the rBDP rate, *β_ν_*(*t*) follows from equation A.2 as below. This constraint means we’ve removed a free parameter.

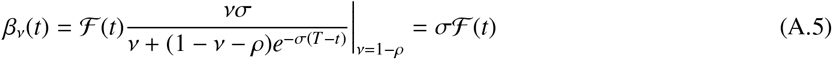

The lack of dependence of the per lineage rBDP rate on *T* or *t* is striking. This sampled BDP therefore yields a reconstructed tree that matches a Yule-Furry process with speciation rate *λ* – *μ*. Consequently all of the analysis from section 3.3 is applicable with *λ* replaced by *σ* in those equations. Furthermore, for these parameter settings the BDP will present an analogous inference problem to that of the Kingman coalescent [18].

As a final point, we comment that there are many other ways to sample (both deterministically and randomly) the extant lineages for the reconstructed process. Unfortunately, these are more involved than homochronous sampling and do not yield simple relationships [5]. However, once the resulting sampled rBDP conforms to a Markov birth process then the SF can be applied. Investigating the application of the SF to various sampling measures will form a key part of our future work.

